# Contrasting resistance and resilience to light variation of the coupled oxic and anoxic components of an experimental microbial ecosystem

**DOI:** 10.1101/2021.08.13.456201

**Authors:** Marcel Suleiman, Frank Pennekamp, Yves Choffat, Owen L. Petchey

## Abstract

Understanding how microbial communities of aquatic ecosystems respond to environmental change remains a critical challenge in microbial ecology. In this study, we used phototrophic oxic-anoxic micro-ecosystems to understand how the functioning and diversity of aerobic and anaerobic lake analog communities is affected by light deprivation. Continuous measurements were performed to describe oxygen dynamics (mean/min/max/amplitude) and time-series of full-length 16S rRNA sequencing were used to quantify changes in alpha- and beta-diversity. In the top oxic layer, oxygen concentration decreased significantly under light deprivation, but showed resilience in mean, minimum and maximum after light conditions were restored. Only the amplitude of diurnal fluctuations in oxygen concentrations did not recover fully, and instead tended to remain lower in treated ecosystems. Alpha-diversity of the top oxic layer communities showed a delayed increase after light conditions were restored, and was not resilient. In contrast, alpha-diversity of the anoxic bottom layer communities increased due to the stressor, but was resilient. Community composition changed significantly during light deprivation, showed resilience in the oxic layer and lack of resilience in the anoxic layer. Alpha-diversity and the amplitude of oxygen within and among treatments were strongly correlated, suggesting that higher diversity could lead to less variable oxygen concentrations, or vice versa. Our experiment showed that light deprivation induces multifaceted responses of community function and structure, hence focusing on a single stability component could potentially be misleading.

## Introduction

Microbial communities are critical components of ecosystems, driving their development and functioning since billions of years. Over these extreme time-scales, microbes have tackled major transitions, such as the increasing oxygenation of Earth’s atmosphere. The success of microbial life is to a large degree explained by being highly adaptable to a variety of environmental conditions [1, 2]. Even slight changes in conditions can lead to taxonomical and functional community shifts, in turn being important indicators of microbial community stability, or lack thereof [3]. Communities can react towards disturbances by showing resistance, resilience, functional redundancy or alternate stable states [4]. Observing responses on the functional and compositional level and how they are related can provide much needed insights about the mechanisms of microbial community stability [5]. How functioning and stability are mediated by the diversity of an ecosystem has been addressed with controlled experiments [6–8], however, these assume random species loss, while disturbances often affect biodiversity non-randomly [9].

Lake ecosystems are facing a broad range of stressors [10], including temperature-increase, fertilizer-intake [11] chemical pollutants [12], or increasing microplastic pollution [13]. One factor that is still not investigated well is light deprivation of lake ecosystems, which can occur indirectly due to biomass formation on aquatic surfaces, including blooms of algae [14] and biofilms[15], as well as plant biomass growth. A broad range of functional aerobic and anerobic microbial groups are highly dependent on light, including cyanobacteria in the upper water column and phototrophic sulfur bacteria in the lower water column. Hence light deprivation could have a large effect on microbial community composition and ecosystem properties such as the oxygen concentration in lake ecosystems [16] due to tight coupling of light, microbial respiration and cyanobacterial O_2_ production.

Here we studied the response of mixed aerobic-anaerobic communities to light deprivation within a single ecosystem with a recently developed dynamic phototrophic oxic-anoxic micro-ecosystem [17]. These micro-ecosystems are analogs of freshwater ecosystems, like lakes and ponds, which harbor highly diverse functional groups of microorganisms, tightly connected through the oxygen state of the aquatic environment, which are dependent on light for oxygenic and anoxygenic photosynthesis.

## Material and methods

Sediment and water samples were taken from a pond (Zurich, Switzerland, 47°23’51.2”N 8°32’33.3”E) and eight micro-ecosystems were set-up as reported previously [17]. In short, 1.5 cm sediment (with 0.5 % crystalline cellulose, 0.5 % methyl-cellulose, 1 % CaSO_4_, 0.2 % CaCO_3_, 0.01 % NH_4_H_2_PO_4_) were covered with 16 mL pond water and incubated for 35 days at 24 °C under a light-dark-cycle of 16:8 h (gradient of light). Continuous non-invasive oxygen measurements (PreSens Precision Sensing GmbH, Germany) were performed every 5 min at the bottom liquid part (1.5 cm above sediment) and top liquid part (2 cm below surface) of each column. After incubating the eight columns for eight days, four columns were covered tightly with aluminum foil and incubated in darkness for seven days (stressor-treatment). After the treatment, incubation continued with the standard light conditions, like the control group, for another 20 days. 500 µL liquid sample were taken on day 8 (prior-to-stressor), day 15 (stressor-sample), day 19 (short-term recovery) and day 35 (long-term recovery), respectively, at the height of the top and bottom oxygen sensor. DNA extraction and 16S rRNA full-length sequencing (PacBio) were performed as reported previously [17]. Raw sequencing data were transcribed to Amplicon Sequence Variants (ASV) with Dada2 [18], and analysis of alpha- and beta-diversity were performed with the R packages phyloseq [19] and vegan [20].

In order to compare the effects of the light treatment we analysed seven response variables (daily mean, maximum, minimum, and amplitude of oxygen concentration, alpha diversity (Shannon index), and two components of microbial community composition). Microbial community composition was quantified using NMDS (non-metric multidimensional scaling) based on Bray-Curtis distances with the *metaMDS* function of the vegan R package [20], with two dimensions used (giving the previously mentioned two components of microbial community composition).

Each of the seven response variables was analysed separately at each time point it was measured at. At each time point, we calculated the mean of each treatment group (control group and light deprivation group), the difference between these means, and the 95% confidence interval of this difference (assuming normally distributed errors). We then visually inspected if the confidence interval included zero, and if so judged there to be no difference between the treatment and control group, and if so that there was a difference. We say that a response variable was resistant to the treatment if the 95% confidence interval of the difference between control and treatment included zero at the end of the treatment (day 15 and 19) (or not resistant if the 95% CI did not include zero). We say that a response variable was resilient to the treatment if the 95% confidence interval of the difference between control and treatment included zero at the end of the experiment (day 35) (or not resilient if the 95% CI did not include zero). Although the oxygen measurements are plotted and analysed for each day, we only based assessments of resistance and resilience on the same days as for community composition (days 15, 19, 35). This reduces the number of comparison, which reduces any issues associated with multiple testing, and also reduces any issues associated with temporal autocorellation / repeated measured. Detailed scripts are available on zenodo (10.5281/zenodo.5195092) and sequencing raw reads are available on NCBI (PRJNA731625).

## Results and discussion

Each of the eight micro-ecosystems showed oxygen-based stratification (Figure 1a) and microbial community compositions characteristic for layered lake ecosystems, with aerobic communities in the oxic top liquid part (diverse members of *Gammaproteobacteria, Bacteroidia, Cyanobacteriia*) and (phototrophic) sulfur bacteria in the anoxic bottom layer (*Chlorobium, Chlorobaculum, Magnetospirillum, Sulfuricurvum*) (Figure 1, Figure 1-figure supplement 1).

**Figure 1.**
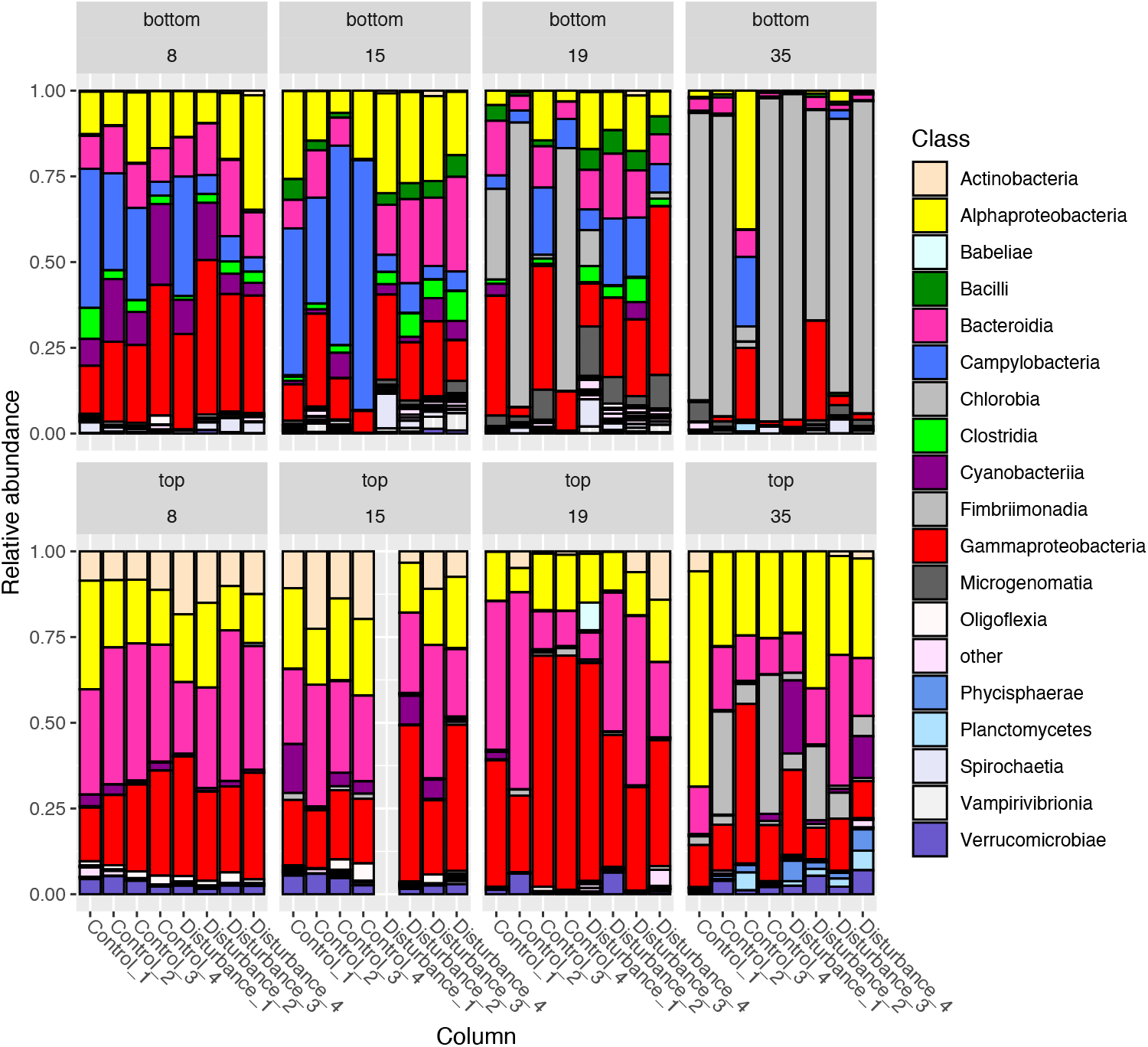
Microbial community composition of the micro-ecosystems. Relative abundance of microbial community composition (rel. abundance > 5 %) on class level for the bottom and top layer communities of the micro-ecosystems. Controls were incubated under a light-dark cycle of 16:8 h for 35 days. Disturbed columns were incubated under light-dark cycle for 8 days (prior-stressor sample), then incubated in darkness until day 15 (stressor sample), before incubating again under a light-dark cycle of 16:8 h (short-term recovery sample on day 19 and long-term recovery sample day 35).

During the first week, the oxygen concentration of the top layer of the eight micro-ecosystems increased during the light- and decreased during dark-phase, resulting in comparable behavior of the mean-, max- and min-oxygen concentration, as well as the amplitude (Figure 2, day 0-8). During light deprivation (day 8-15) the disturbed columns showed a significant decrease in total oxygen concentration to microaerophilic conditions (Figure 2 b, day 8-15). After normal light conditions were restored, oxygen concentration increased, showing resilience until the end of the experiment for mean, minimum and maximum, but not for the amplitude, which was higher in the controls (Figure 2c). Therefore, light deprivation had a lasting impact on the oxygen amplitude, which stayed within a narrower range of values for communities that experienced the stressor, while untreated ones were more variable. A single replicate showed a higher oxygen concentration at the end of the experiment. Exclusion of that microcosm affected the estimated differences in oxygen concentration (i.e., resistance and resilience), but did not qualitatively change the results. Exclusion also had no effect on the conclusions of the analyses of microbial community composition. Since the bottom layer turned completely anoxic within 2 days, we did not detect any effect of light deprivation on the oxygen pattern there.

**Figure 2.**
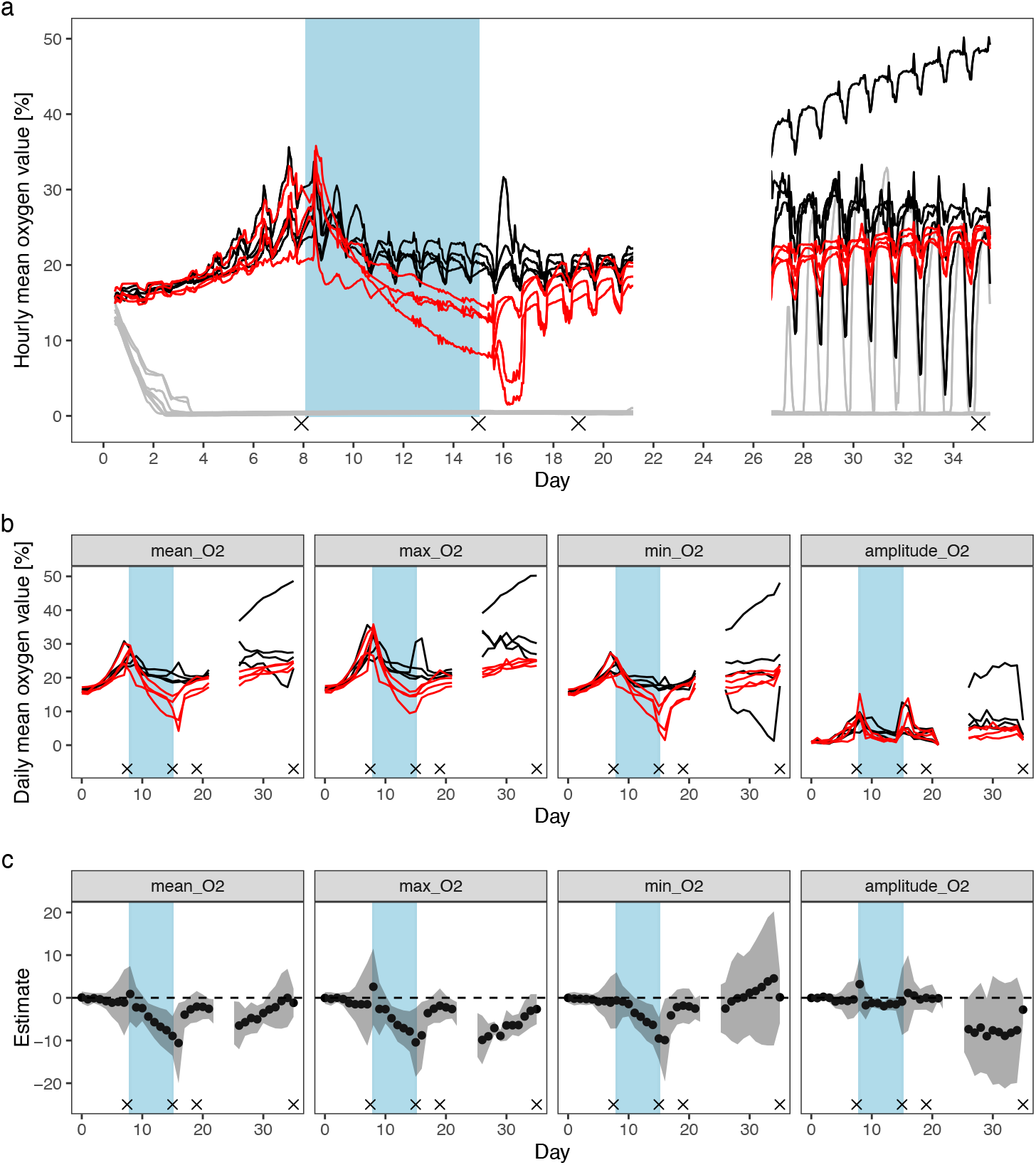
Dynamics and functional response of oxygen in the micro-ecosystems. Oxygen concentration was recorded every 5 minutes. **(a)** Hourly mean of the oxygen concentration of the top sensors (line) and bottom sensors (grey) of the eight micro-ecosystems. **(b)** Daily-mean oxygen concentration (illustrated as mean, maximum, minimum and amplitudes) of the top sensors of the eight micro-ecosystems. **(c)** Estimates of difference in the mean and 95% CI of the estimate of the oxygen concentration treatment vs. controls. Black lines represent controls, red lines represent columns incubated in darkness from day 8-15 (blue area). Gray ribbons show the 95% confidence intervals. Crosses show the sampling days for analysing microbial communities of the top and bottom part. A recording error caused the missing oxygen data from day 23-27.

Alpha diversity was differentially affected by light deprivation in the top and bottom level (Figure 3): The richness of the anaerobic bottom communities was immediately increased by light deprivation (Day 15, Figure 3a+b) and stayed significantly different after normal light conditions were restored in the short-term recovery sample, but showed resilience (i.e. return to control values) in the long-term. In contrast, the alpha-diversity of the aerobic top layer communities was not immediately affected by the stressor, showed marginally significant differences in the short-term recovery samples but stayed significantly changed in the long-term (Figure 3c+d).

**Figure 3.**
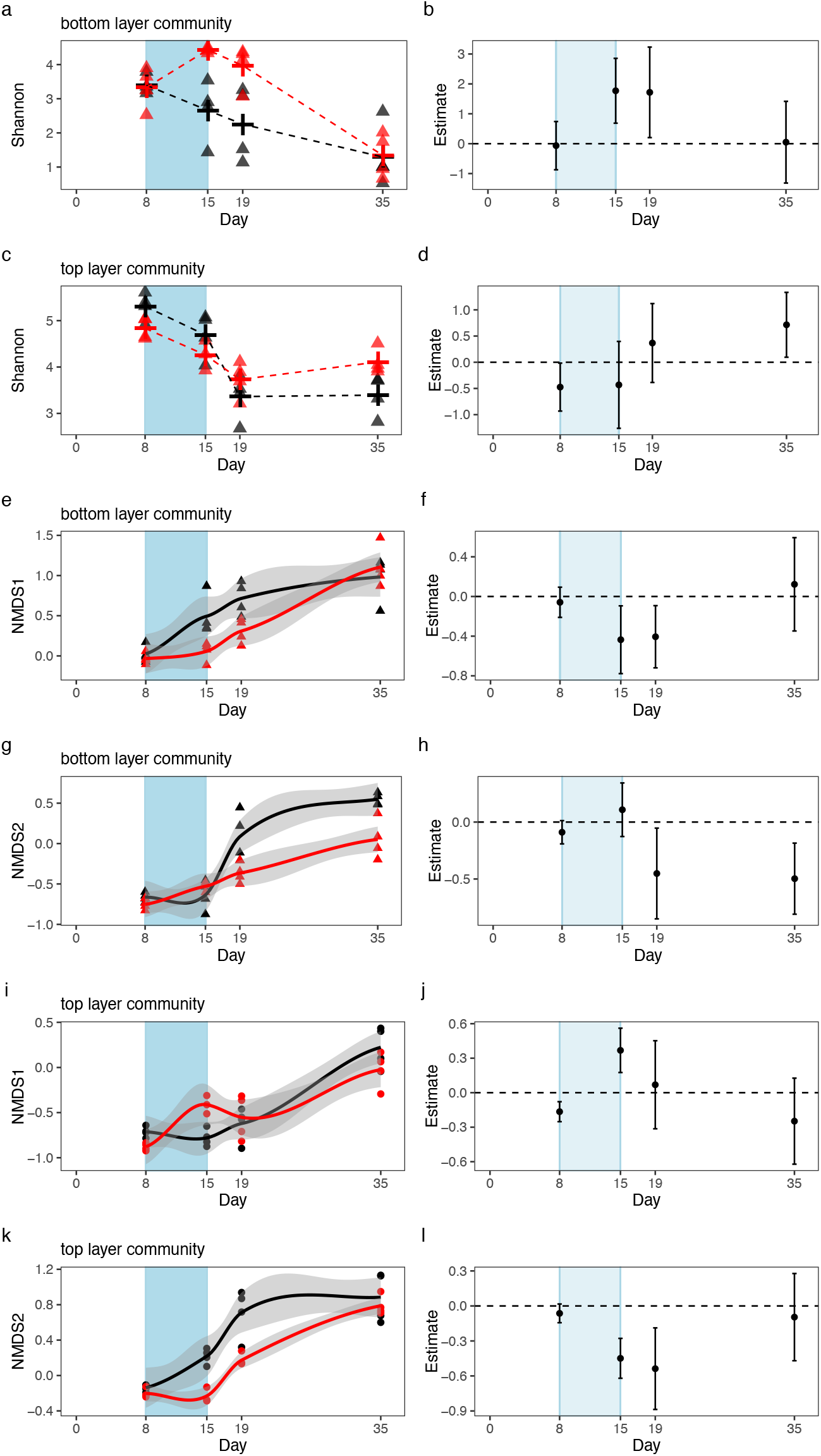
Resistance and resilience of the microbial communities based on full-length 16S RNA sequencing. **(a)** Alpha-diversity based on Shannon index of the bottom water communities of the eight micro-ecosystems. Cross: mean. **(b)** Estimated difference between treatments and 95% confidence interval for the alpha diversity analysis for the bottom water communities. **(c)** Alpha-diversity based on Shannon index of the top water communities of the eight micro-ecosystems. Cross: mean. **(d)** Estimated difference between treatments and 95% confidence interval for the alpha diversity analysis for the top water communities. **(e**,**g**,**i**,**k)** Beta-diversity analysis based on NMDS1 and NMDS2 score for the bottom and top water communities, respectively. **(f**,**h**,**j**,**l)** Estimated difference between treatments and 95% confidence interval for the beta diversity (based on NMDS1 and NMDS2 scores) as response parameters based on NMDS1 score for the bottom and top water communities. Black lines represent controls, red lines represent columns incubated in darkness from day 8-15 (blue area). Gray ribbons show the 95% confidence intervals.

Non-metric multidimensional scaling revealed significantly different composition of top and bottom layer communities within a single micro-ecosystem, but the top and bottom composition converged during light deprivation (Figure 3-figure supplement 1). This convergence is most likely due to the loss of the light- and oxygen-gradient during the disturbance, indicating how stressors can directly influence microbial communities by affecting stratification processes in an ecosystem.

To understand the dynamic changes of community composition, we analysed time series of NMDS1 and NMDS2 (Figure 3b). For the bottom community, the NMDS1 score was immediately affected by light deprivation but showed resilience, while NMDS2 was affected in delayed fashion and remained significantly changed (Figure 3e-h). The top layer communities were affected immediately based on NMDS1 and NMDS2, but both showed resilience at the last sampling (Figure 3i-l).

In order to understand possible relationships between diversity and stability, we examined the correlation of alpha diversity and the oxygen amplitude of the last incubation day 35 (Figure 4). We found a strong negative correlation between alpha-diversity and amplitude of diurnal oxygen concentration (n=8, t= -5.1134, p = 0.002). This correlation holds when excluding the replicate with high mean oxygen concentration, very high amplitude of oxygen variation (∼ 7.4), and low diversity (SI 2.8) (n=7, t= -2.6, p = 0.04) (Figure 4-figure supplement 1). This finding is in line with the insurance hypothesis of biodiversity, that higher biodiversity will cause lower temporal variation in aggregate properties of communities and of ecosystem states, such as oxygen concentration [6]. However, in our experiment diversity was not manipulated, but was rather an outcome of the factors such as environmental conditions (including the light treatment) and interspecific interactions. Accordingly, we cannot be sure if the higher diversity observed in the light deprivation treatment was the cause of the greater stability (lower amplitude) of the oxygen concentration in these communities. Also, since, we did not manipulate the amplitude of the oxygen fluctuations, we cannot say if this was responsible for the observed differences in alpha diversity. Further studies manipulating the diversity of organisms in the ecosystem and separately the magnitude of fluctuations in environmental conditions such as oxygen concentration would be needed to assign causation and understand the likely feedback between organismal diversity and environmental fluctuations.

**Figure 4.**
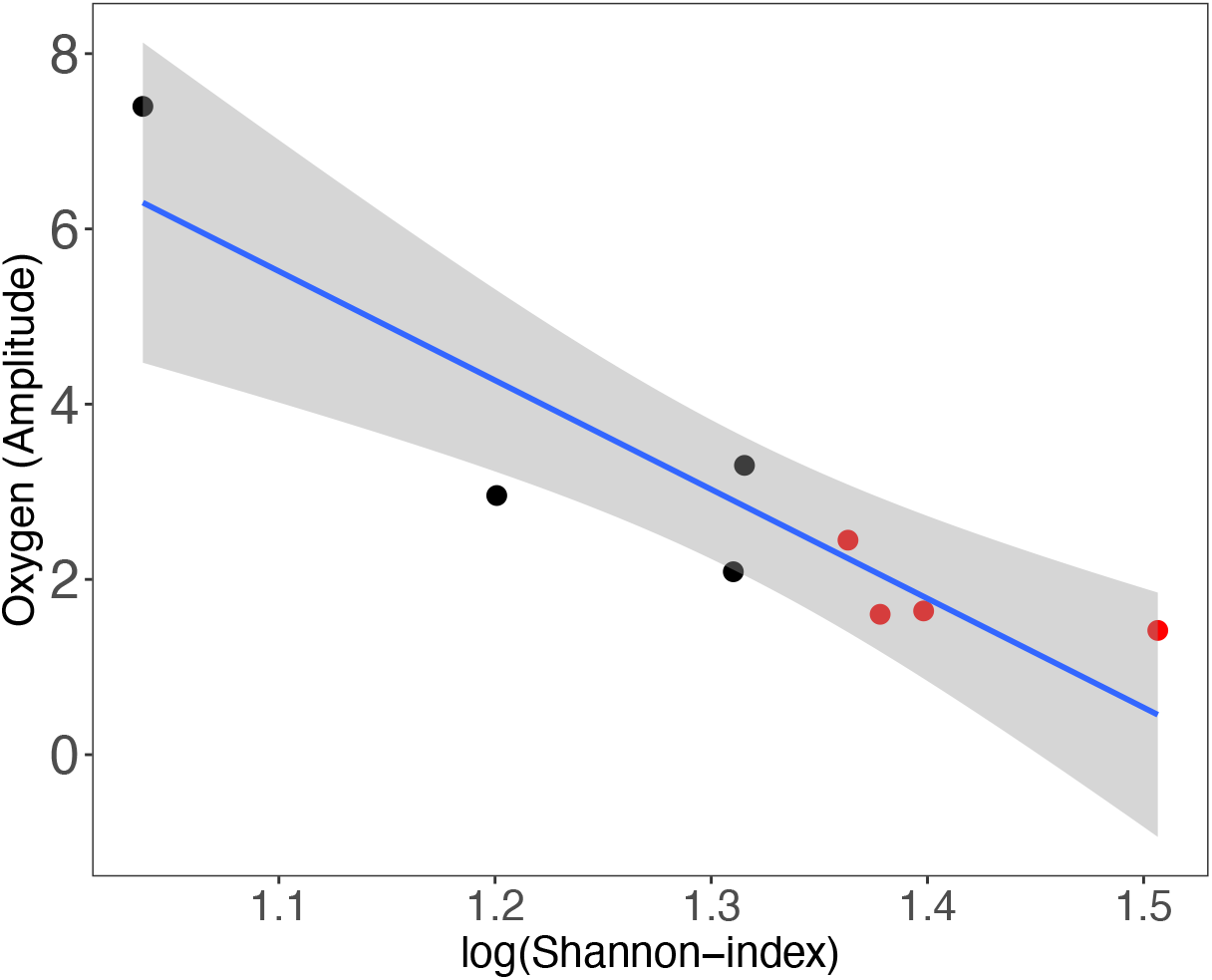
Relationship between the amplitude of daily fluctuations in oxygen concentration and Shannon diversity of top layer communities the long-term recovery sample (day 35). Black symbols represent controls, red symbols represent treated columns. Gray ribbons show the 95% confidence intervals.: Pearson’s product-moment correlation: p-value=0.002, correlation coefficients= -0.90.

Our experiment showed that light deprivation induces multifaceted responses of community function and structure at multiple temporal scales. We found that light deprivation has differential effects on the coupled communities in ecosystems: (i) light deprivation had a strong effect on microbial function, richness and composition, but most responses were resilient in the long run (ii); For non-resilient aspects, we found that the richness and the amplitude of oxygen concentration were related. We suggest that non-random effects of light deprivation leads to a change in richness, with subsequently affects the oxygen amplitude, but this proposed causal relationship needs experimental testing (iii); recovery and resilience operate at different time scales and hence several time points of analysis are needed for making robust conclusions about microbial resistance and resilience [21]. Focusing on just a single aspect of community stability may hence be misleading, since a stressor can affect multiple stability components differently [22, 23].

## Acknowledgements

MS was funded by Forschungskredit of the UZH (FK-20-125). OLP and FP were supported by the University of Zurich Research Priority Programs in Global Change and Biodiversity. OLP was supported by Swiss National Science Foundation (Project 310030_188431). We thank Martin Kapun for bioinformatical support with raw data. We thank the Functional Genomic Center Zurich for sequencing efforts and support with sequencing analysis. We would like to thank Prof. Meredith Schuman for the very helpful feedback on the manuscript.

## Competing interests

The authors declare no competing financial interests.

## Author contributions

OLP and MS planned the experimental set-up. MS performed all experiments in the lab. MS performed the up- and downstream bioinformatics. FP supported the data analysis. YC provided technical support. OLP, MS and FP drafted the manuscript. All authors confirmed the final version of the manuscript.

---

**Figure 1-figure supplement 1.**
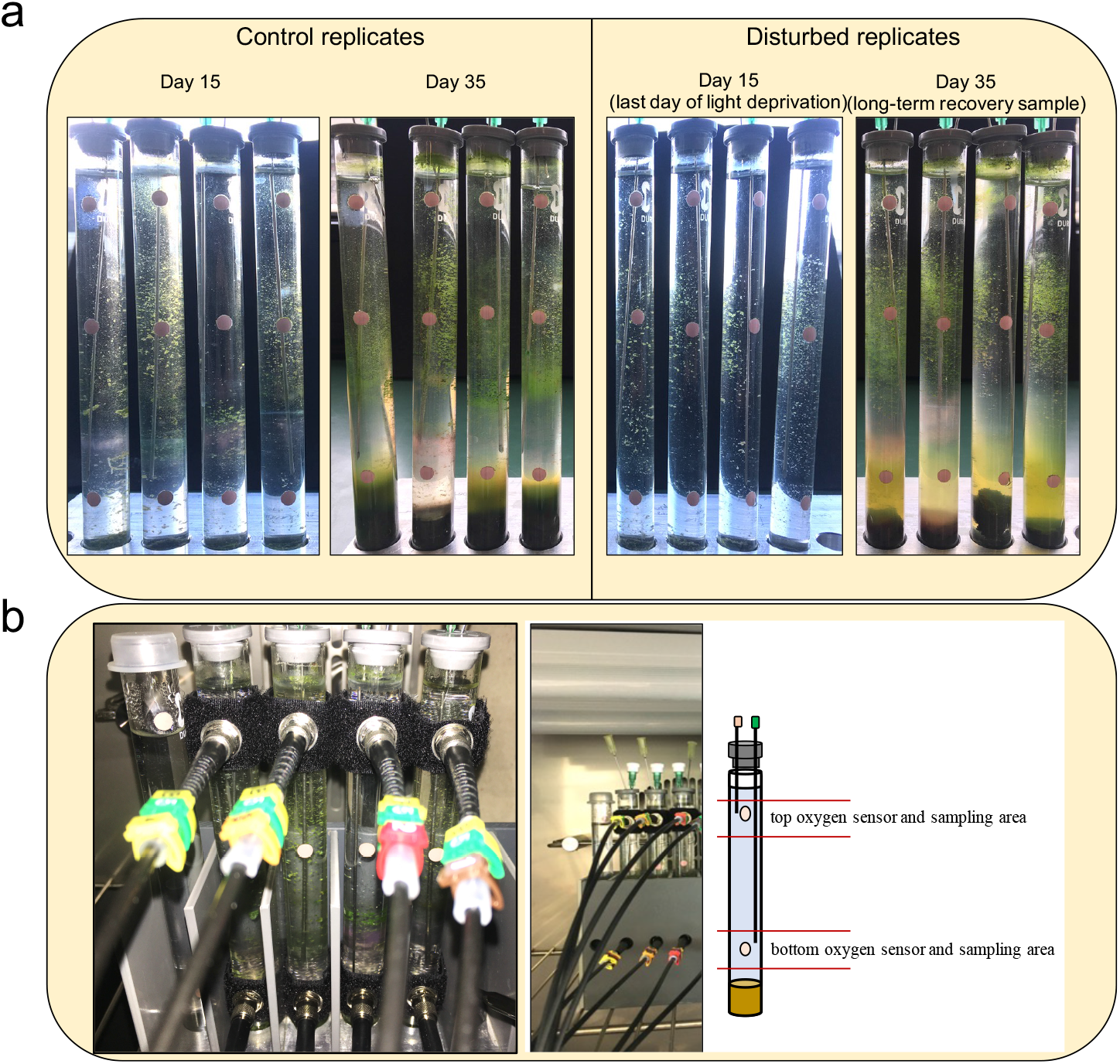
Macroscopic appearance and experimental setup of the micro-ecosystems. Control micro-ecosystems on day 15 and day 35. Disturbed micro-ecosystems on day 15 (stressor sample) and day 35 (long term recovery sample). (b) Technical setup (exemplary cutout) of the micro-ecosystems and the automatic oxygen-measurements at the top and bottom sensors. Oxygen was measured in 5 min intervals. Samples were taken at day 8 (prior stressor sample), day 15 (stressor sample), day 19 (short-term recovery sample) and day 35 (long-term recovery sample) at height of the top and bottom sensor, respectively.

**Figure 3-figure supplement 1.**
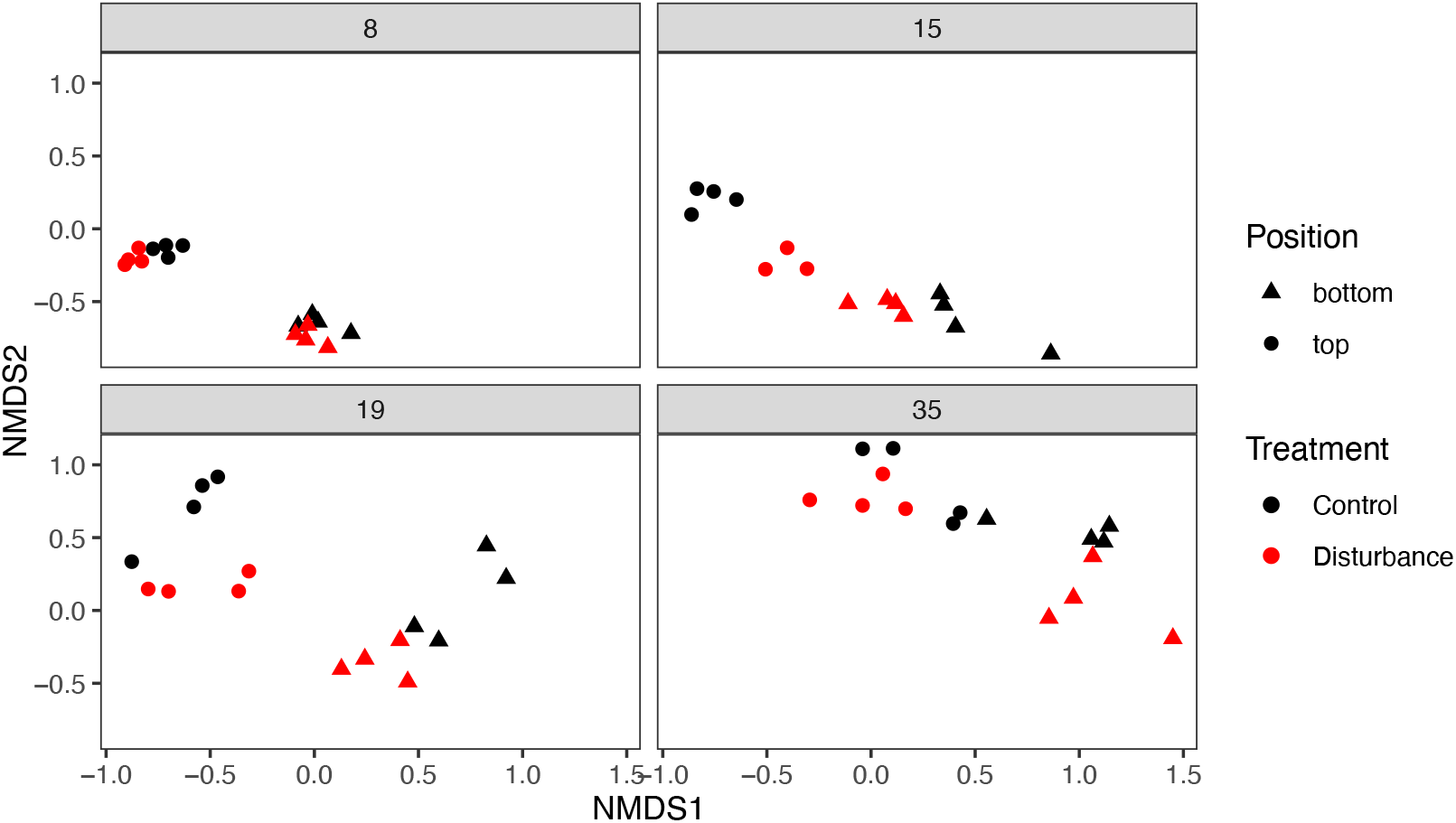
NMDS analysis of the top and bottom communities of the prior-stressor sample (day 8), stressor-sample (day 15), short-term recovery sample (day 19) and long-term recovery sample (day 35). Stress is 0.16.

**Figure 4-figure supplement 1.**
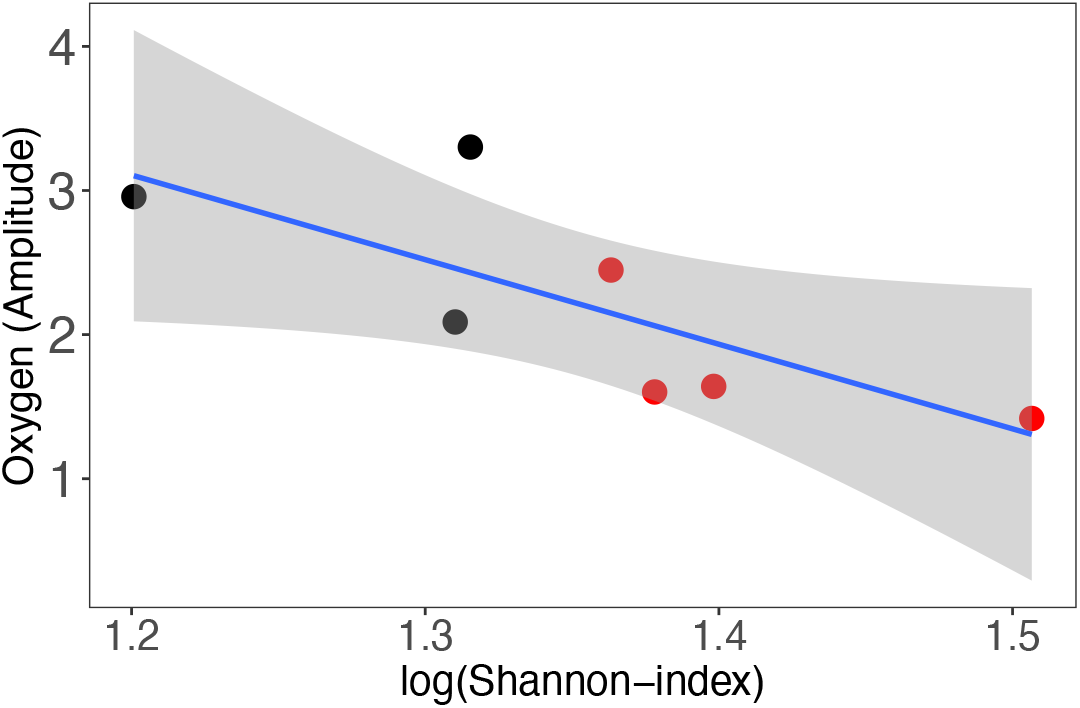
Relationship between the amplitude of daily fluctuations in oxygen concentration and Shannon diversity of top layer communities without the high amplitude-low diversity replicate in the long-term recovery sample (day 35). Black symbols represent controls, red symbols represent treated columns. Gray ribbons show the 95% confidence intervals. Pearson’s product-moment correlation: p-value=0.047, correlation coefficients = -0.76.

## Notes

### Competing Interest Statement

The authors have declared no competing interest.

